# Effects of short-term resistance training and tapering on maximal strength, peak power, throwing ball velocity, and sprint performance in handball players

**DOI:** 10.1101/586586

**Authors:** Souhail Hermassi, Aloui Ghaith, René Schwesig, Roy J. Shephard, Mohamed Souhaiel Chelly

**Affiliations:** Sport Science Program, College of Arts and Sciences, Qatar University, Doha, Qatar; Research Unit (UR17JS01) Sport Performance, Health & Society, Higher Institute of Sport and Physical Education, Ksar-Saîd, University of “La Manouba”, Tunis, Tunisia; Department of Orthopaedic and Trauma Surgery, Martin-Luther-University Halle-Wittenberg, Halle, Germany; Faculty of Kinesiology and Physical Education, University of Toronto, Toronto, Canada; Department of Biological Sciences Applied for Physical Activities and Sport, Higher Institute of Sport and Physical Education of Ksar Said, University of “La Manouba”, Tunis, Tunisia

**Keywords:** Peak power, throwing velocity, vertical jump, sprint performance, maximal force, periodization

## Abstract

The purpose of this study was to assess the effect of short-term resistance training and two weeks of tapering on physical performances in handball players. Following a ten-week progressive resistance training program, subjects were divided between an experimental (n = 10) and a control group (n = 10). The experimental group completed a resistance training program, followed by a two-week period when the training intensity was tapered by 60%, while the control group maintained their typical pattern of training. Muscle power (force–velocity test and squat and counter-movement jump tests), sprinting ability (10m and 30m), ability to change direction (T-half test) and throwing velocity (a 3-step throw with a run, and a jump throw) were evaluated before training, at the end of training and after tapering. The experimental group showed significantly larger interaction effects for the 10-week training period (12/15, 80%), than for the following 2 weeks of tapering (10/15, 67%), with the largest gains being in 15 m sprint times (d=3.78) and maximal muscular strength in the snatch (d=3.48). Although the performance of the experimental group generally continued to increase over tapering, the mean effect size for the training period was markedly higher (d=1.92, range: 0.95-3.78) than that seen during tapering (d=1.02, range:−0.17-2.09). Nevertheless the ten weeks of progressive resistance training followed by two weeks of tapering was an effective overall tactic to increase muscle power, sprint performance and ball throwing velocity in handball players.

## INTRODUCTION

The concept of a tapering of training was first introduced by Costill et al. [1] for competitive swimmers. The efficacy of tapering has subsequently been well documented in runners [2], swimmers [3], cyclists [4], rowers [5], and triathletes [6], with improvement in performance or its physiological correlates [7–10]. From a neuromuscular perspective, tapering usually increases muscular strength and power, often with associated gains in performance at the muscular and whole body level. Oxidative enzyme activities can also increase, along with positive changes in single muscle fibre size, metabolic properties and contractile properties.

The aim of tapering is to reduce the physiological and psychological stress imposed by heavy daily training and thus to optimize competitive performance. It can be carried out in many ways, including either progressive or stepped reductions in training volume, intensity, and frequency [2]. The length of the taper period has also varied widely [7–11]. Several studies have shown that a 2-week taper period [12–14] provides significant improvements in performance, while others have reported improvements over relatively short periods (<7 days) [4] and much longer periods (>28 days) [9]. Some studies have reduced training volume by as much as 85% [15], whereas others have shown similar improvements in performance after only a 31% decrease [14].

No previous study has investigated the effects of a two-week step-tapering period on the physical performance characteristics of handball players undergoing resistance training. The objective of this study was to thus analyze the effects of a two-week step tapering on upper and lower-limb muscle power, ball throwing velocity, jump performance and sprinting ability in elite male handball players. We hypothesized that players who completed two weeks of tapering would show increases in performance relative to those seen at the end of the initial training period.

## MATERIALS AND METHODS

### Participants

All procedures were approved by the Institutional Review Committee for the ethical use of human subjects, according to current national and international laws and governing regulations. Participants gave their written informed consent after receiving both a verbal and a written explanation of the experimental design and its potential risks. Subjects were free to withdraw from the study without penalty at any time. A questionnaire covering medical history, age, height, body mass, training characteristics, injury history, handball experience, and performance level was completed before participation. An initial examination by the team physician focused on orthopedic and other conditions that might preclude resistance training; however, all participants were found to be in good health.

All participants were drawn from the First National League, with playing positions as follows: (pivots, n=3; backs, n=4; wings, n=4; CG: pivots, n=1; backs, n=4; wings, n=4.). All playing positions were included, since every activity has a special feature, based on playing position. Five players were left-handed. Participants were randomly divided between experimental and control groups. These groups were initially well matched in terms of physical characteristics (experimental group: age: 20.9 ± 0.7 years, body mass: 85.2 ± 8.8 kg, height: 1.84 ± 0.03 m, body fat 13.7 ± 0.8%; control group: age: 20.6 ± 0.5 years, body mass: 85.6 ± 9.4 kg, height: 1.82 ± 0.04 m, body fat 13.7 ± 0.6%).

### Experimental Design

This study used a pre-test post-test design. A group of 20 male handball players volunteered for random assignment to either a weightlifting training + tapering group (Experimental group) (n = 10) or a control group that continued to follow the standard in-season regimen (n =10). Both groups had been training for 5 months, and were already 4 months into the competitive season before introduction of the modified training program. All completed two familiarization trials in the 2 weeks before definitive testing, which was carried out before training (T_0_), after 10 weeks of added weightlifting (T_1_) and after 2 weeks of tapering only in the experimental group (T_2_).

Assessments included sprint times over 5m, 15m and 30m, throwing velocity, vertical jump, and the strength and power of both the upper and lower limbs. Testing sessions were carried out at the same time of the day, and under the same experimental conditions, at least 3 days after the most recent competition. Players maintained their normal intake of food and fluids, but abstained from physical exercise for 1 day before testing, drank no caffeine-containing beverages for 4 hours before testing, and ate no food for 2 hours before testing. Strong verbal encouragement ensured maximal effort throughout both measurement and resistance training sessions.

#### Procedures

Strength training portion was introduced over a 10-week period (January to April) from the 22^nd^ to the 29^th^ week of the playing season, immediately after the traditional 8-day winter holiday. Both the experimental and control groups were accustomed to moderate strength training (1 session per week of bench press and half squat exercises at 60-80% of 1-RM loading). All had also participated in the standard handball training program since the beginning of the competitive season. This routine consisted of six 90 minute training sessions per week, plus a competitive game played on the weekend. Physical conditioning was performed three times per week; it aimed at the development of strength, and incorporated elements of high-intensity interval training, weight-lifting, plyometric, power lifting and gymnastics. Anaerobic training was based on plyometric and sprint training drills, and aerobic fitness was developed using small-sided games. Training sessions consisted mainly of technical-tactical skill development (60% of session time) and strength and conditioning routines (40% of session time). During the 10 week intervention, the control group maintained their standard training while the experimental group replaced a part of their regimen (technical-tactical skill development) with a resistance training program twice per week (Figure 1).

**Figure 1.**
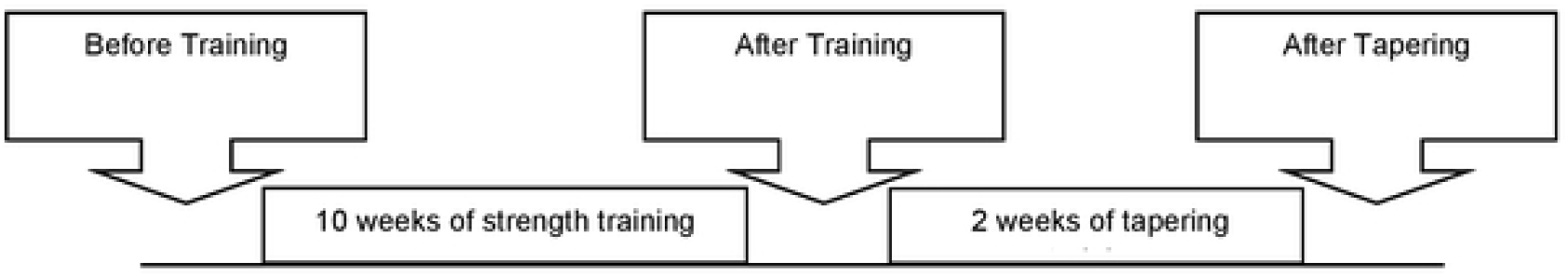
Study design.

#### Testing Schedule

All tests were performed on the same indoor handball court, under similar ambient conditions (temperature, 22.1±0.5 C; relative humidity 60±5%), and at the same time of day (5:00 p.m. to 7:00 p.m.). Intensive training was avoided for 24 h, and participants also fasted for three hours prior to testing. Standardized warm-up exercises preceded all maximal efforts; these included 5 min of low intensity running, 3 x 30-m progressive accelerations, and a maximal 30-m sprint, interspersed with 3-minute periods of passive recovery. The warm-up prior to throwing tests included push-ups with both hands on the ground, 8 to 10 free-ball throws, and exercises such as trunk rotation, trunk side-bends, trunk wood-chops, and internal and external rotational movements of the shoulder, with the arm held at 90° shoulder abduction and 90° elbow flexion to simulate the throwing position.

Testing was integrated into the participants’ weekly training schedules. Familiarization two weeks before definitive testing determined individual 1-repetition max (1-RM) values for the different strength tests. The three definitive assessments were made before training, after 10 weeks of added resistance training and after 2 weeks of tapering. All sets of tests were administered on three non-consecutive days, using the same procedures and technicians who were blinded to group assignment. On the first day, anthropometric assessments were followed by squat countermovement jumps and finally force–velocity testing of first the upper and then the lower limbs. On the second day, sprint performance was assessed, followed by maximal repetition bench press (1-RM_BP_) and maximal repetition snatch (1-RM_snatch_). On the third day, ball throwing velocity, maximal repetition clean (1-RM_clean_) and maximal repetition jerk (1-RM_jerk_) were determined

### Day one

#### Anthropometry

Standard equation equations were used to predict the percentage of body fat from the biceps, triceps, subscapular, and suprailiac skinfold readings [16]:

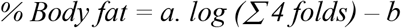

where Σ S is the sum of the four skinfold readings (in mm), and *a* and *b* are constants dependent on sex and age.

#### Squat Jump and Countermovement Jump

Characteristics of the squat jump and the countermovement jump (jump height, maximal force before take-off, maximal velocity before take-off and the average power of the jump) were determined using a force platform (Quattro Jump, version 1.04, Kistler Instrument AG, Winterthur, Switzerland). Jump height was determined as the center of mass displacement, calculated from the recorded force and body mass. Subjects began the SJ with theirs hips and knees in 90 degrees of flexion and performed a vertical jump by pushing upwards and ballistically, extending their hips and knee, and keeping their legs straight throughout. The countermovement jump began with the subjects in an upright position, making a rapid downward movement to approximately 90 degrees of hip and knee flexion then reversing this motion by ballistically moving into full extension. One minute of rest was allowed between the three total trials of each exercise, and the best score was recorded.

#### The Force–Velocity Test

Force–velocity measurements for the lower limbs were performed on a standard Monark cycle ergometer (model 894 E, Monark Exercise AB, Vansbro, Sweden) [17]. In brief, the instantaneous maximal pedaling velocity during a 7-second all-out sprint was used to calculate the maximal anaerobic power for each braking force, and the subject was judged to have reached peak power (Wpeak) if an additional load induced a decrease in power output. Arm tests were made using an appropriately modified cycle ergometer [17,18]. The parameters measured included Wpeak, and the maximal braking force maximal pedaling velocity for upper limbs [17,18]. Arm tests began with a braking force equal to 1.5% of the subject’s body mass [17,18]. After a 5-minute recovery, the braking force was increased in a sequence of 2, 3, 4, 5, 6, 7, 8, and 9% of body mass [17,18].

### Day two

#### 30-m Sprint Performance

The 30m sprint began with a standardized warm-up. Subjects then ran 40m, with times at 5, 15, and 30m recorded by a series of paired photocells (Microgate, Bolzano, Italy). Three trials were separated by 6-8 min of recovery. Subjects began from a standing position, with the front foot 0.2 m behind the starting photocell beam.

#### 1-RM Bench Press

The maximal strength of the upper extremity was assessed using a maximum one repetition successive eccentric-concentric bench press action (1-RM_BP_). The bench press (elbow extension) was chosen because it involves arm muscles such as the triceps and the pectorals that are specific to overhand throwing. The test was performed in a squatting apparatus; the barbell was attached at both ends, and linear bearings on two vertical bars allowed only vertical movements. The bar was initially positioned 10 mm above the subject’s chest and supported by the bottom stops of the measuring device. The participant was instructed to perform an eccentric contraction followed by a concentric contraction from the starting position, maintaining the shoulders at 90-degree abduction throughout the test. No bouncing or arching of the back was allowed. A warm-up was provided and consisted of five repetitions at 40–60% of each subjects’ perceived maximum. Thereafter, four to five separate attempts with 2 min rest intervals were performed until the person was unable to extend the arms fully. The last acceptable extension was recorded as the 1-RM_BP_.

#### One Repetition Maximum Snatch

Subjects were required to lift the loaded barbell upwards using a wide grip from the starting position of hip and knee flexion with one movement until both arms were locked in an extended position. Thereafter the lifter moved from a low squat to a standing position. After some warmup lifts with lighter weights the following lifts were performed: 2 at 70%, 2 at 80%, 1 at 90%, 1 at 95%, and 2-3 at 100% of the one repetition maximum (1-RM) [19].

### Day three

#### Handball Throwing

Explosive strength was evaluated on an indoor handball court via a 3-step running throw and a jump throw. After a 10 minute standardized warm-up, participants threw a standard handball (mass 480 g, circumference 58 cm). They were allowed to put resin on their hands before throwing with maximal velocity towards the upper right corner of the goal. The coaches supervised these tests closely to ensure that the appropriate techniques were followed. Each individual continued until three correct throws had been recorded, up to a maximum of three sets of three consecutive throws. A 1-to 2-minute rest was allowed between sets and 10–15 seconds between two throws in the same set. For the jump throw, players made a preparatory three-step run before jumping vertically and releasing the ball while in the air, behind a line 9 m from the goal. For the running throw, players took a preparatory run limited to three regular steps before releasing the ball, behind the line, 9 m from the goal. Throwing time was recorded with an accuracy of 1ms, using a digital video camera (HVR to A1U DV Camcorder; Sony, Tokyo, Japan). The camera was positioned on a tripod 2 m above and perpendicular to the plane of ball release. Data processing software (Regavi & Regressi, Micrelec, Coulommiers, France) converted measures of handball displacement to velocities. The validity of the camera and the data processing software under working conditions was verified [20] by measuring the speed of rolling balls (2–14 m/s) by camera (Vc) and checking data over a given distance (3 m) against photoelectric cells (Vpc) (GLOBUSREHAB and Sports High Tech, Articolo ERGO TIMER, Codognè, Italy). The 2 estimates of speed were well correlated (Vc = 0.9936Vpc + 0.65; r=0.99; p<0.0001) [20]. The throw with the greatest average velocity was selected for further analysis.

#### One Repetition Maximum Clean and Jerk

In the Olympic clean and jerk, the loaded barbell was lifted in a single movement with a shoulder wide grip and the knees bent from the starting position on the platform to the chest. The participant stood from this low squat position, and thereafter lifted the loaded barbell by extending their arms to bring the barbell overhead [19].

#### T-half test

The T-half test [21] was performed using the same protocol as the t-test, except that the total distance covered was reduced from 36.56 to 20 m and inter-cone distances were modified. Criteria for acceptable test trials were the same as in the t-test, with recording of the better of two final trials (test–retest session). Subjects began from a standing position, with the front foot 0.2 m behind the starting photocell beam (Microgate, Bolzano, Italy).

#### 1-RM Back Half-Squat

Participants maintained an upright position throughout. The bar was grasped firmly with both hands and was also supported on the subject’s shoulders. The hips and knees were initially bent to 90 degrees and were then fully extended during the concentric portion of the test. Warm-up consisted of a set of five repetitions at loads 40%–60% of the perceived maximum. To measure the 1-RM, the barbell was loaded with free weights to an initial 90% of the pretest 1-RM. Two consecutive tests were made, and if the 2 repetitions were mastered, a load of 5 kg was added after a recovery interval of 3 minutes. When the participant had performed 2 successful repetitions of his pretest RM value, further loads of 1 kg were added after the recovery interval [22]. If the second repetition could not be completed with the new loading, the corresponding load was considered as the individual’s 1-RM (Hermassi et al., 2011).

#### Weightlifting training program

All training sessions were supervised by certified strength and conditioning specialists knowledgeable in weightlifting guidelines and pedagogy. Participants were encouraged to increase the amount of weight lifted and to achieve concentric fatigue within each designated repetition range, and all completed a minimum of 95% of scheduled sessions. A standardized warm-up including jogging, dynamic stretching exercises, calisthenics, and fundamental weightlifting exercises specific to their training program; further, sessions ended with 5 minutes of cool-down activities that included dynamic stretching.

Each training program session comprised four different exercises of 3 sets of 5–10 repetitions [22]; all were compound lifts involving multi-articular movements and multiple muscle groups. Volume and intensity of effort were based on previous recommendations for handball players [23], and the ability of each individual [24] as established from his 10 repetition maximum in the selected resistance exercises. Participants were required to lift their maximum weight for a given number of repetitions while using proper technique. Instructors reviewed technique and made appropriate adjustments in loading during each training session. If the required number of repetitions could not be completed within a set, individuals were given 30 seconds to 1 minute of rest before attempting to complete the set again.

The training program comprised four Olympic-style exercises: snatch from a squatting position, bench press, half-squat, and clean and jerk, using a certified weightlifting bar with Olympic plates. During the first 2 weeks, 3-4 sets of 6-8 repetitions of each exercise were performed. The initial load corresponded to 60% of 1-RM, and 3 minutes of rest was allowed between sets. During the third, fourth, and fifth weeks, the volume was increased to 3 sets of 6-10 repetitions, with loads corresponding to 70% of 1-RM respectively. For the final 5 weeks, 3 sets of 5-10 reps at 75–85% of 1-RM were performed. During weekends 11 and 12, the training volume was decreased by approximately two-thirds and training frequency by 50% (Table 1).

**Table 1.**
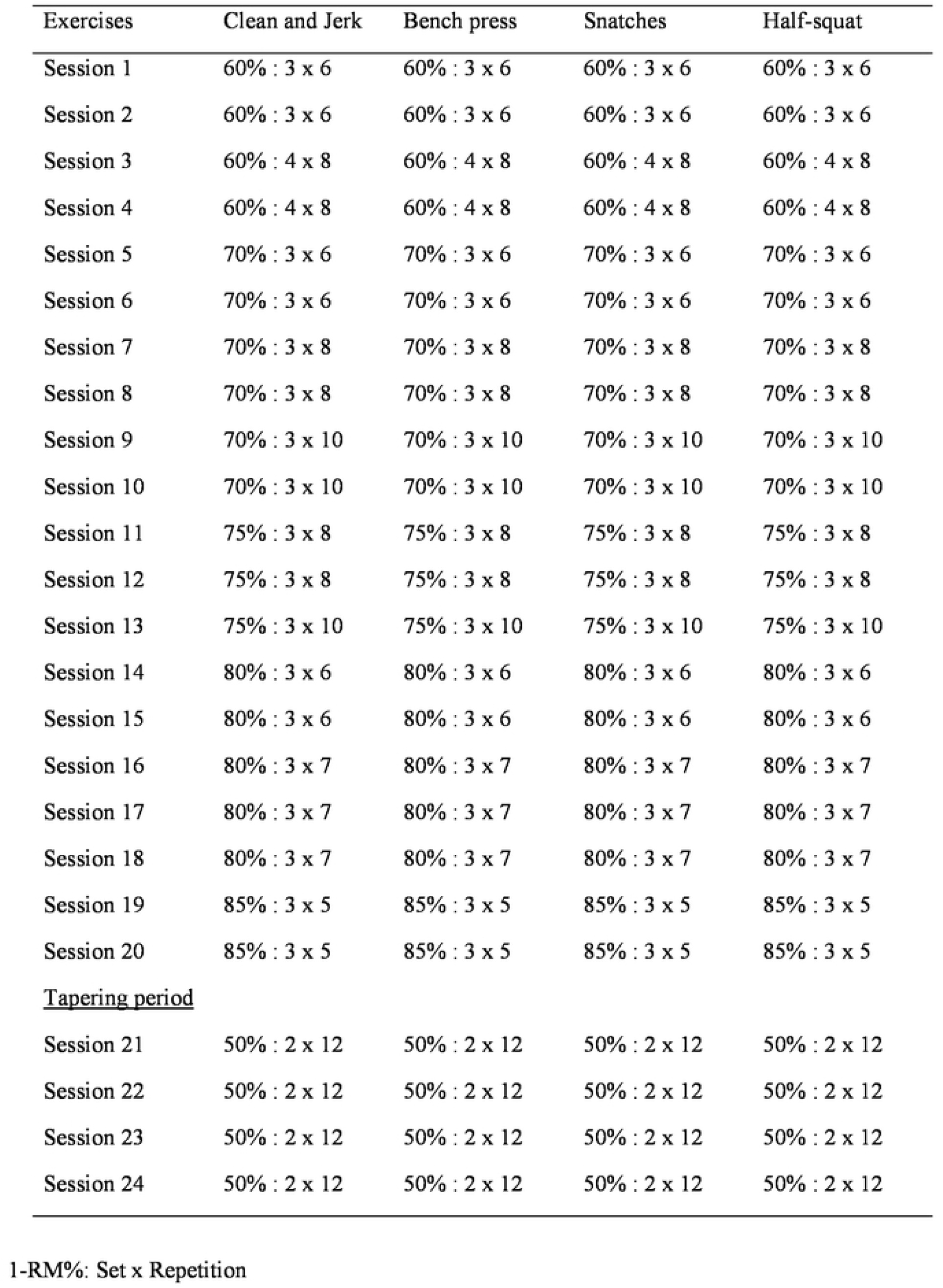
Weight-lifting training program and tapering period as followed over 12 weekends.

| Exercises | Clean and Jerk | Bench press | Snatches | Half-squat |
| --- | --- | --- | --- | --- |
| Session 1 | 60% : 3 x 6 | 60% : 3 x 6 | 60% : 3 x 6 | 60% : 3 x 6 |
| Session 2 | 60% : 3 x 6 | 60% : 3 x 6 | 60% : 3 x 6 | 60% : 3 x 6 |
| Session 3 | 60% : 4 x 8 | 60% : 4 x 8 | 60% : 4 x 8 | 60% : 4 x 8 |
| Session 4 | 60% : 4 x 8 | 60% : 4 x 8 | 60% : 4 x 8 | 60% : 4 x 8 |
| Session 5 | 70% : 3 x 6 | 70% : 3 x 6 | 70% : 3 x 6 | 70% : 3 x 6 |
| Session 6 | 70% : 3 x 6 | 70% : 3 x 6 | 70% : 3 x 6 | 70% : 3 x 6 |
| Session 7 | 70% : 3 x 8 | 70% : 3 x 8 | 70% : 3 x 8 | 70% : 3 x 8 |
| Session 8 | 70% : 3 x 8 | 70% : 3 x 8 | 70% : 3 x 8 | 70% : 3 x 8 |
| Session 9 | 70% : 3 x 10 | 70% : 3 x 10 | 70% : 3 x 10 | 70% : 3 x 10 |
| Session 10 | 70% : 3 x 10 | 70% : 3 x 10 | 70% : 3 x 10 | 70% : 3 x 10 |
| Session 11 | 75% : 3 x 8 | 75% : 3 x 8 | 75% : 3 x 8 | 75% : 3 x 8 |
| Session 12 | 75% : 3 x 8 | 75% : 3 x 8 | 75% : 3 x 8 | 75% : 3 x 8 |
| Session 13 | 75% : 3 x 10 | 75% : 3 x 10 | 75% : 3 x 10 | 75% : 3 x 10 |
| Session 14 | 80% : 3 x 6 | 80% : 3 x 6 | 80% : 3 x 6 | 80% : 3 x 6 |
| Session 15 | 80% : 3 x 6 | 80% : 3 x 6 | 80% : 3 x 6 | 80% : 3 x 6 |
| Session 16 | 80% : 3 x 7 | 80% : 3 x 7 | 80% : 3 x 7 | 80% : 3 x 7 |
| Session 17 | 80% : 3 x 7 | 80% : 3 x 7 | 80% : 3 x 7 | 80% : 3 x 7 |
| Session 18 | 80% : 3 x 7 | 80% : 3 x 7 | 80% : 3 x 7 | 80% : 3 x 7 |
| Session 19 | 85% : 3 x 5 | 85% : 3 x 5 | 85% : 3 x 5 | 85% : 3 x 5 |
| Session 20 | 85% : 3 x 5 | 85% : 3 x 5 | 85% : 3 x 5 | 85% : 3 x 5 |
| <u>Tapering period</u> |  |  |  |  |
| Session 21 | 50% : 2 x 12 | 50% : 2 x 12 | 50% : 2 x 12 | 50% : 2 x 12 |
| Session 22 | 50% : 2 x 12 | 50% : 2 x 12 | 50% : 2 x 12 | 50% : 2 x 12 |
| Session 23 | 50% : 2 x 12 | 50% : 2 x 12 | 50% : 2 x 12 | 50% : 2 x 12 |
| Session 24 | 50% : 2 x 12 | 50% : 2 x 12 | 50% : 2 x 12 | 50% : 2 x 12 |

1-RM%: Set x Repetition

#### Statistical Analyses

All statistical analyses were performed using SPSS version 25.0 for Windows (SPSS Inc., Chicago, IL, USA). Means and standard deviations of dependent variables were calculated across participants. Differences between groups (experimental vs. control) and sessions (pre-vs. post-intervention) were tested using a two-factor (time, group) univariate general linear model [25] for both phases of the intervention (session 1 vs. 2: effects of program training; session 2 vs. 3: effects of tapering period). Percentage changes were calculated as ([posttraining value - pre-training value]/pre-training value) x 100. Comparisons between initial and final tests used non-paired Student t-tests. The effect size (d) (mean difference of scores divided by the pooled standard deviation) was calculated for each parameter [26]. After applying a Bonferroni correction (p<0.05 divided by the number of tests (14)), the significance level (p) was set with p<0.003. Consequently, differences between means (group, time and group-time effects) were considered as being statistically significant if: p<0.003 and η^2^>0.20 and d≥0.5 [25]. Due to the relatively small number of cases in each group (n=10), decisions on significance were based on all three statistical values in order to avoid an overestimation of intervention. Applying a power calculation for this study design and assuming that p<0.01, 1-β=0.80 and d=0.5, 80 subjects per group would have been necessary in order to test the hypothesis conclusively [25].

## RESULTS

Compliance of the experimental group with the added training was high, each exercise session being completed with a high level of motivation and effort. The data demonstrated significant interaction effects (group x time) in 12 of 15 parameters for the 10-week intervention (Table 2, figure 2 figure 3 figure 4 figure 5). The effect sizes were all larger than 0.90; the greatest gain was in 15 m sprint time (d=3.78), and the largest interaction effect was for the half squat (p<0.001, η^2^=0.827). Three parameters (30 m sprint, squat jump height, and the relative power of lower limb) showed no significant interaction effects. Two relevant (d≥−0.5) decreases of performance (absolute power of lower limb: d=−0.86; snatch: d=−0.66) were seen in the control group (Table 2).

**Table 2.** Comparison between the experimental and control groups before and after 10-weck training period (examination 1 vs. 2). Significant interaction effects and effect sizes arc highlighted in bold.

| Parameter | Experimental Group<br>(Mean±SD) |  |  | Control Group<br>(Mean±SD) |  |  | Variance analysis/ effects |  |  |  |  |  |
| --- | --- | --- | --- | --- | --- | --- | --- | --- | --- | --- | --- | --- |
|  | Pre | Post |  | Pre | Post |  | Group |  | Time |  | Group x time |  |
|  |  |  | d |  |  | d | p | η<sup>2</sup> | p | η<sup>2</sup> | p | η<sup>2</sup> |
| Sprint performance and ability to change direction (s) |  |  |  |  |  |  |  |  |  |  |  |  |
| Sprint 15 m | 2.74±0.08 | 2.40±0.10 | **3.78** | 2.96±0.62 | 3.11±0.56 | 0.03 | 0.020 | 0.265 | 0.054 | 0.191 | **<0.001** | **0.601** |
| Sprint 30 m | 4.69±0.18 | 4.41±0.20 | 0.95 | 4.68±0.24 | 4.77±0.27 | 0.09 | 0.066 | 0.176 | 0.036 | 0.222 | **0.001** | **0.493** |
| Agility t-half test | 6.86±0.46 | 6.02±0.36 | **2.05** | 6.34±0.24 | 6.35±0.34 | -0.04 | 0.525 | 0.023 | **<0.001** | 0.620 | **<0.001** | **0.639** |
| Throwing performance (m/s) |  |  |  |  |  |  |  |  |  |  |  |  |
| Jump shot | 21.9±3.1 | 25.6±3.7 | **1.07** | 25.7±4.1 | 26.0±2.8 | 0.09 | 0.177 | 0.099 | **<0.001** | 0.500 | **0.001** | **0.437** |
| Running shot | 27.7±2.2 | 32.4±2.1 | **2.19** | 25.4±3.8 | 24.9±3.6 | -0.14 | 0.001 | 0.442 | **<0.001** | 0.629 | **<0.001** | **0.730** |
| Jump performance (cm) |  |  |  |  |  |  |  |  |  |  |  |  |
| CMJ | 39.1±2.2 | 42.2±1.6 | **1.63** | 40.3±2.0 | 40.6±2.1 | 0.15 | 0.773 | 0.005 | **<0.001** | 0.670 | **<0.001** | **0.573** |
| SJ | 39.8±2.3 | 42.2±1.8 | **1.18** | 38.6±2.3 | 39.4±2.2 | 0.36 | 0.039 | 0.216 | 0.001 | 0.490 | 0.039 | 0.215 |
| 1RM (kg) |  |  |  |  |  |  |  |  |  |  |  |  |
| half squat | 159±12 | 177±12 | **1.46** | 136±17 | 129±19 | -0.39 | **<0.001** | 0.605 | 0.001 | 0.491 | **<0.001** | **0.827** |
| bench press | 74.5±7.6 | 86.0±9.9 | **1.31** | 81.7±10.8 | 78.1±9.3 | -0.36 | 0.933 | 0.000 | 0.003 | 0.406 | **<0.001** | **0.713** |
| snatch | 55.5±4.4 | 68.9±3.3 | **3.48** | 61.5±5.8 | 57.5±6.3 | -0.66 | 0.167 | 0.104 | 0.002 | 0.420 | **<0.001** | **0.713** |
| clean and jerk | 63.0±5.9 | 73.8±5.4 | **1.92** | 64.0±8.1 | 63.5±7.8 | -0.06 | 0.125 | 0.126 | **<0.001** | 0.558 | **<0.001** | **0.603** |
| **Power of lower limb** |  |  |  |  |  |  |  |  |  |  |  |  |
| Power (W) | 845±116 | 995±85 | **1.38** | 836±85 | 747±118 | -0.86 | 0.008 | 0.333 | 0.064 | 0.178 | **<0.001** | **0.762** |
| Power (W/kg) | 10.0±1.6 | 11.8±1.5 | **1.69** | 9.9±1.3 | 8.9±1.7 | 0.35 | 0.035 | 0.224 | 0.078 | 0.162 | **<0.001** | **0.752** |
| **Power of upper limb** |  |  |  |  |  |  |  |  |  |  |  |  |
| Power (W) | 438±54 | 486±30 | **3.18** | 454±64 | 431±58 | -0.38 | 0.372 | 0.044 | 0.276 | 0.065 | **0.004** | **0.377** |
| Power (W/kg) | 5.1±0.8 | 5.7±0.5 | **0.92** | 5.7±1.1 | 5.4±0.9 | -0.20 | 0.710 | 0.008 | 0.390 | 0.041 | **0.008** | **0.332** |

**CMJ = countermovement jump; SJ = squat jump.**

**Figure 2:**
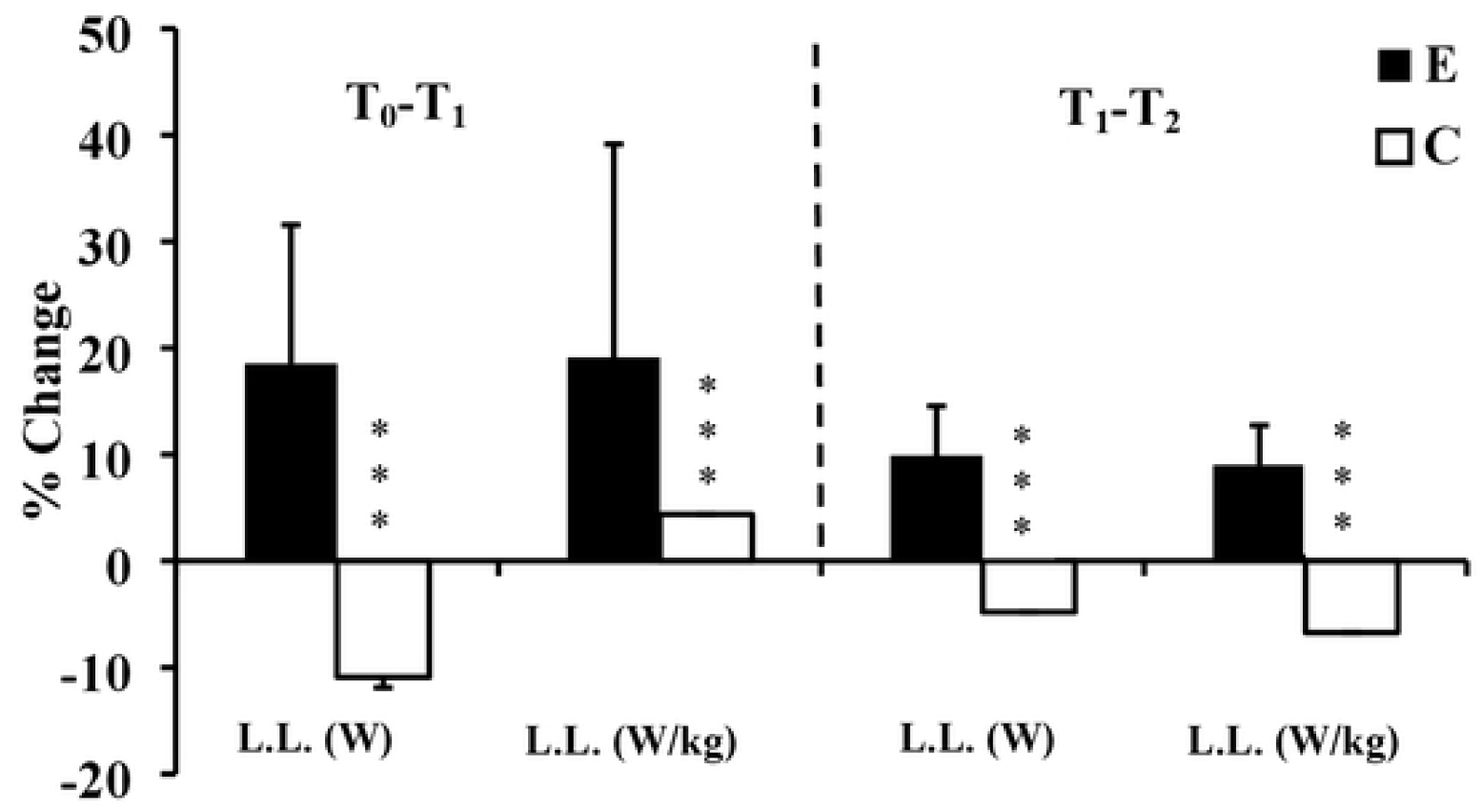
Percentage changes of upper limb power at T_1_, and T_2_ for Experimental (E) and Control (C) groups. T_0_: before training; T_1_: after 10 weeks of resistance training; T_2_: after 2 weeks of tapering; L.L: lower limb; ***: ANOVA group x time interaction significantly different between E and C at the level of *p* < 0.001.

**Figure 3:**
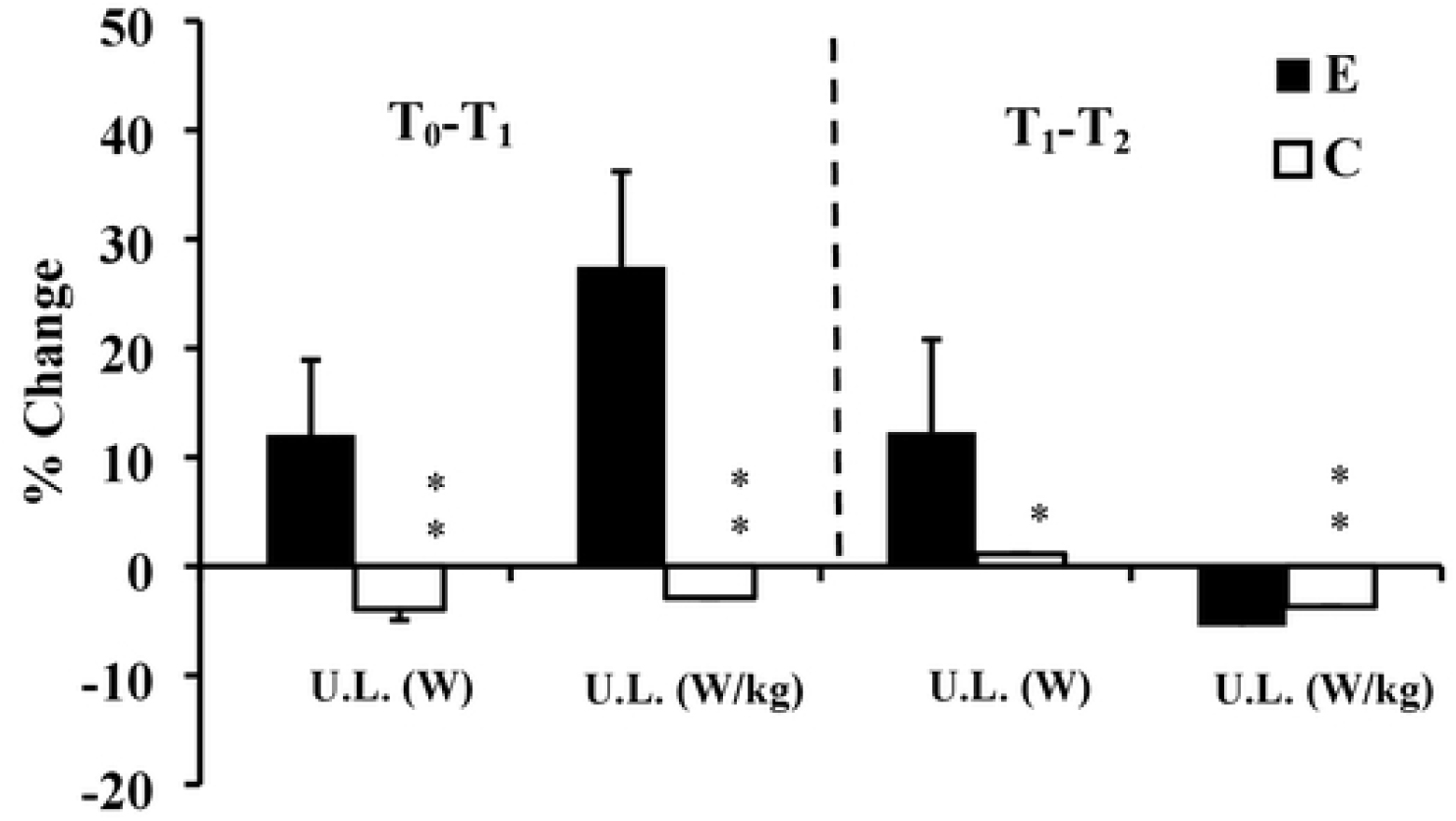
Percentage changes of power of upper limb at T_1_, and T_2_ for Experimental (E) and Control (C) groups. T_0_: before training; T_1_: after 10 weeks of resistance training; T_2_: after 2 weeks of tapering; U.L: upper limb; *: ANOVA group x time interaction significantly different between E and C at the level of *p* < 0.05; **: ANOVA group x time interaction significantly different between E and C at the level of *p* < 0.01.

**Figure 4:**
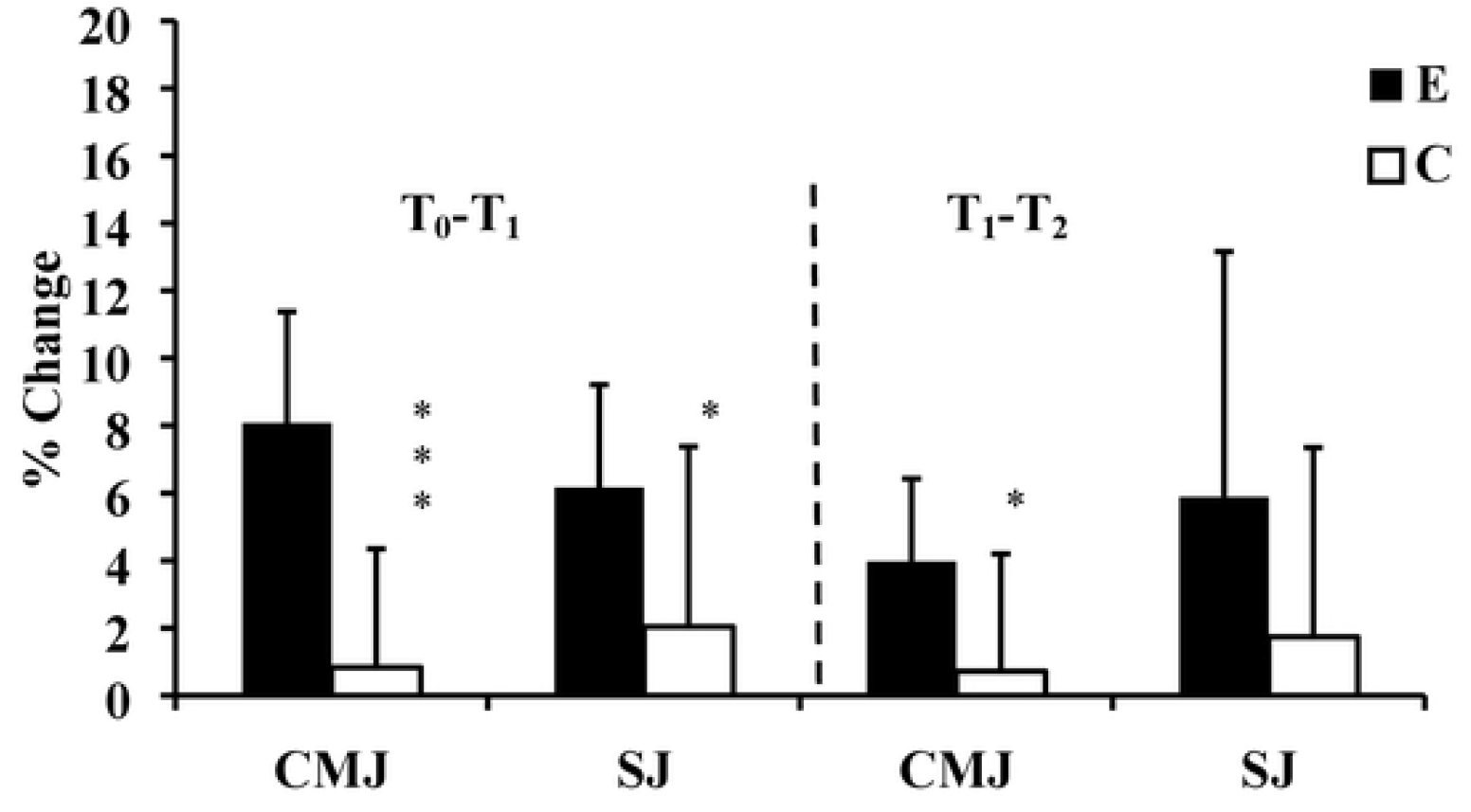
Percentage changes of vertical jump at T_1_, and T_2_ for Experimental (E) and Control (C) groups. T_0_: before training; T_1_: afer 10 weeks of resistance training; T_2_: after 2 weeks of tapering; CMJ: Counter-movement Jump; SJ: Squat Jump; *: ANOVA group x time interaction significantly different between E and C at the level of *p* < 0.05; ***: ANOVA group x time interaction significantly different between E and C at the level of *p* < 0.001.

**Figure 5:**
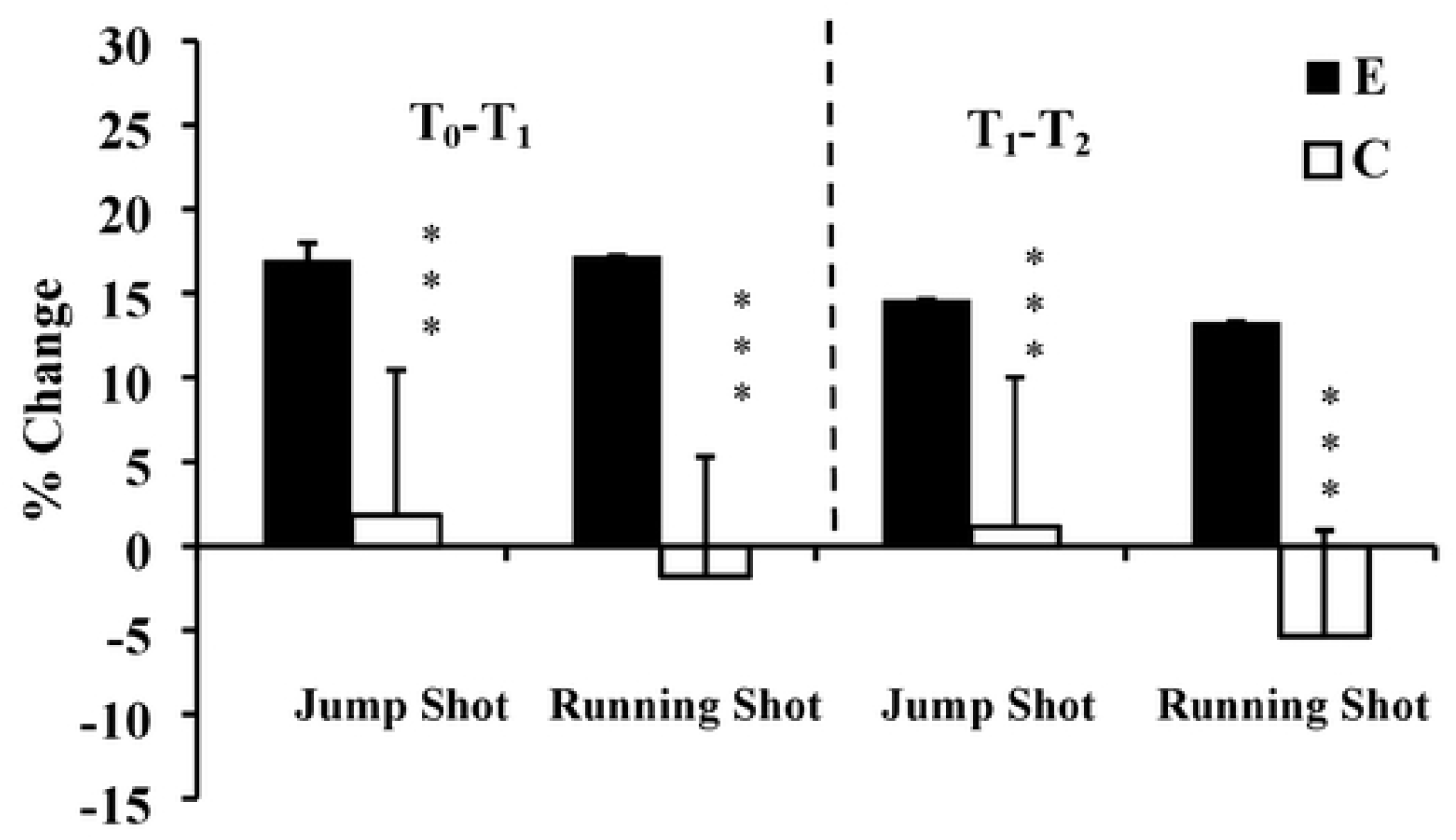
Percentage change of t ball throwing velocity at T_1_, and T_2_ for Experimental (E) and Control (C) groups. To: before training; T_1_: after 10 weeks of resistance training; T_2_: after 2 weeks of tapering; ***: ANOVA group x time interaction significantly different between E and C at the level of *p* < 0.001.

During the 2-week tapering period, changes were smaller (Table 3), but 10 of 15 parameters showed further interaction effects.

**Table 3.** Comparison between experimental and control groups before and after the 2-week tapering period. Significant interaction effects and both effect sizes arc highlighted in bold.

| Parameter | Experimental Group<br>(Mean±SD) |  |  | Control Group<br>(Mean±SD) |  |  | Variance analysis/ effects |  |  |  |  |  |
| --- | --- | --- | --- | --- | --- | --- | --- | --- | --- | --- | --- | --- |
|  | Pre | Post |  | Pre | Post |  | Group |  | Time |  | Group x time |  |
|  |  |  | d |  |  | d | p | η² | p | η² | p | η² |
| Sprint performance and ability to change direction (s) |  |  |  |  |  |  |  |  |  |  |  |  |
| Sprint 15 m | 2.40±0.10 | 2.16±0.13 | **2.09** | 3.11±0.56 | 3.37±0.55 | -0.01 | <0.001 | 0.646 | 0.843 | 0.002 | **<0.001** | **0.451** |
| Sprint 30 m | 4.41±0.20 | 4.40±0.12 | 0.06 | 4.77±0.27 | 4.87±0.17 | -1.03 | <0.001 | 0.630 | 0.311 | 0.057 | 0.230 | 0.079 |
| Agility t-half test | 6.02±0.36 | 6.06±0.11 | -0.17 | 6.35±0.34 | 6.55±0.28 | -0.65 | 0.002 | 0.409 | 0.054 | 0.190 | 0.207 | 0.087 |
| Throwing performance (m/s) |  |  |  |  |  |  |  |  |  |  |  |  |
| Jump shot | 25.6±3.7 | 29.4±4.8 | **0.89** | 26.0±2.8 | 26.2±3.1 | 0.07 | 0.390 | 0.041 | <0.001 | 0.519 | **0.001** | **0.457** |
| Running shot | 32.4±2.1 | 36.7±2.6 | **1.84** | 24.9±3.6 | 23.6±4.4 | -0.33 | <0.001 | 0.739 | <0.001 | 0.596 | **<0.001** | **0.829** |
| Jump performance (cm) |  |  |  |  |  |  |  |  |  |  |  |  |
| CMJ height | 42.2±1.6 | 43.9±1.5 | 1.10 | 40.6±2.1 | 40.9±2.9 | 0.12 | 0.018 | 0.272 | 0.002 | 0.428 | 0.023 | 0.255 |
| SJ height | 42.2±1.8 | 44.7±3.8 | 0.90 | 39.4±2.2 | 40.1±3.7 | 0.24 | 0.006 | 0.346 | 0.015 | 0.288 | 0.158 | 0.108 |
| 1RM (kg) |  |  |  |  |  |  |  |  |  |  |  |  |
| half-squat | 177±12 | 188±10.0 | 0.99 | 129±19 | 131±28 | 0.09 | <0.001 | 0.704 | 0.016 | 0.282 | 0.094 | 0.148 |
| bench press | 86.0±9.9 | 93.5±9.1 | **0.79** | 78.1±9.3 | 74.5±8.9 | -0.40 | 0.004 | 0.372 | 0.010 | 0.313 | **<0.001** | **0.785** |
| snatch | 68.9±3.3 | 74.0±2.1 | **1.88** | 57.5±6.3 | 56.0±6.1 | -0.24 | <0.001 | 0.734 | 0.005 | 0.364 | **<0.001** | **0.658** |
| clean and jerk | 73.8±5.4 | 79.5±4.9 | **1.11** | 63.5±7.8 | 63.0±7.9 | -0.06 | <0.001 | 0.541 | 0.001 | 0.480 | **<0.001** | **0.567** |
| **Power of lower limb** |  |  |  |  |  |  |  |  |  |  |  |  |
| Power (W) | 995±85 | 1086±77 | **1.11** | 747±118 | 698±113 | -0.42 | <0.001 | 0.752 | 0.097 | 0.145 | **<0.001** | **0.659** |
| Power (W/kg) | 11.8±1.5 | 12.8±1.5 | 0.15 | 8.9±1.7 | 8.3±1.6 | -0.36 | <0.001 | 0.617 | 0.060 | 0.183 | **<0.001** | **0.666** |
| **Power of upper limb** |  |  |  |  |  |  |  |  |  |  |  |  |
| Power (W) | 486±30 | 545±23 | **1.62** | 431±58 | 444±72 | -0.12 | 0.001 | 0.457 | 0.002 | 0.428 | 0.032 | 0.232 |
| Power (W/kg) | 5.7±0.5 | 6.4±0.5 | **0.89** | 5.4±0.9 | 5.2±0.6 | -0.20 | 0.007 | 0.336 | 0.095 | 0.147 | **0.009** | **0.322** |

**CMJ = countermovement jump; SJ = squat jump.**

The largest interaction effect: p<0.001, η^2^=0.829) was for the running throw, and the largest further gain was for the 15 m sprint (d=2.09). On the other hand, we noted a reduction in scores for the agility t-half test (d=−0.17). Scores continued to increase during the tapering period, although based on effect sizes (Hartmann et al., 1992), increases were generally smaller than during the 10 week resistance training period (∅d_TG1_=1.92vs. ∅d_TG2_=1.02).

The performance of the control group generally remained stable over the period when the experimental group were tapering, with some trend to regression of performance in a few items, particularly the 30 m sprint test showed a marked decline over the tapering period (d=−1.03, Table 3).

## DISCUSSION

The present data show substantial gains in many measures of performance over the two-week period of tapering. The percentage gains in peak power of the lower limbs and squat jump performance (12.1% and 4% respectively) are lower than seen in some previous trials such as de Lacey et al [27] (who found increases of 45% and 35% in jump height and maximal power respectively, Table 4). However the improvement of countermovement jump performance (4%) is similar to the 5% increase of this same jump seen by Pritchard et al [28] (Table 4).

**Table 4.**
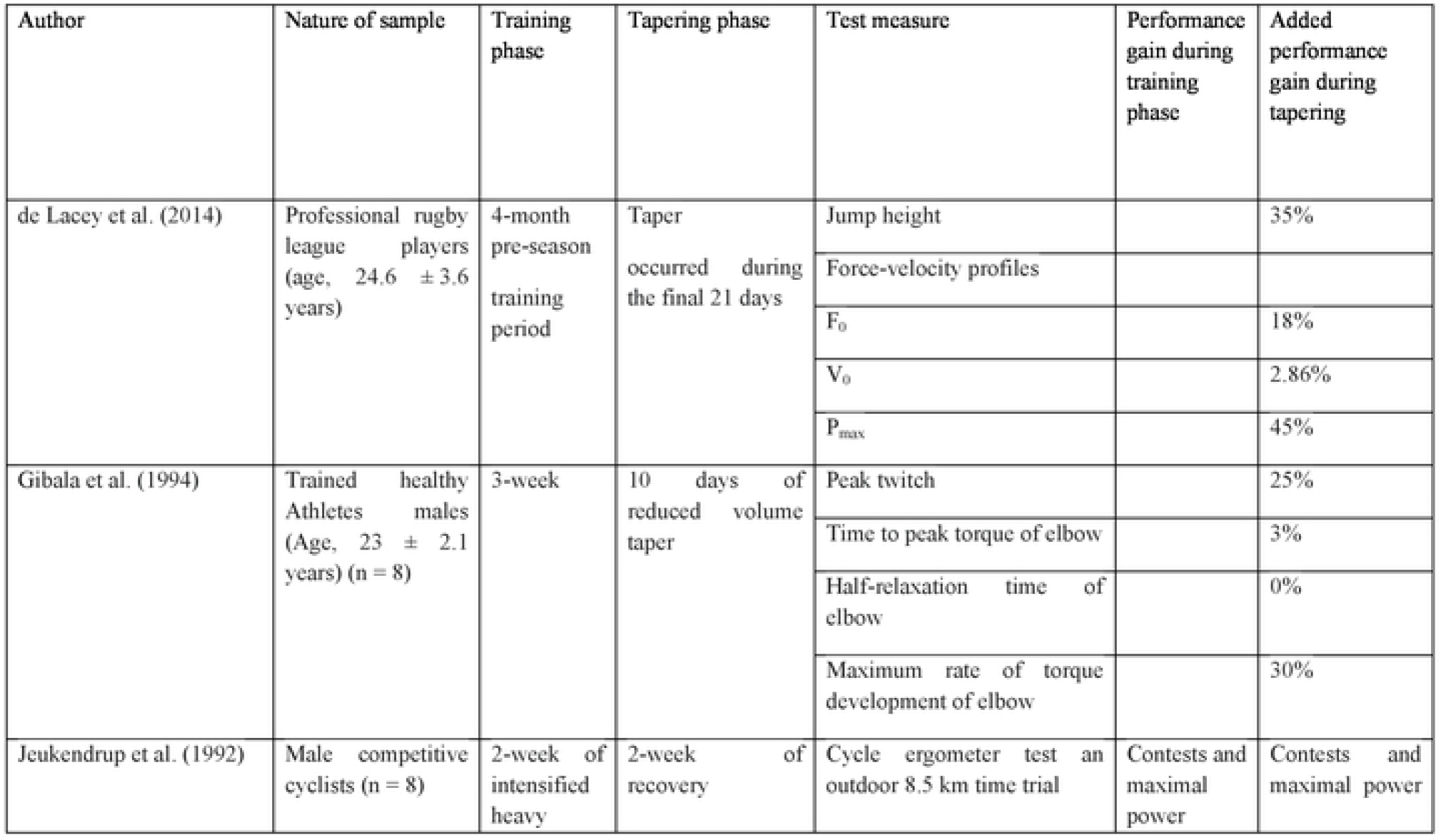
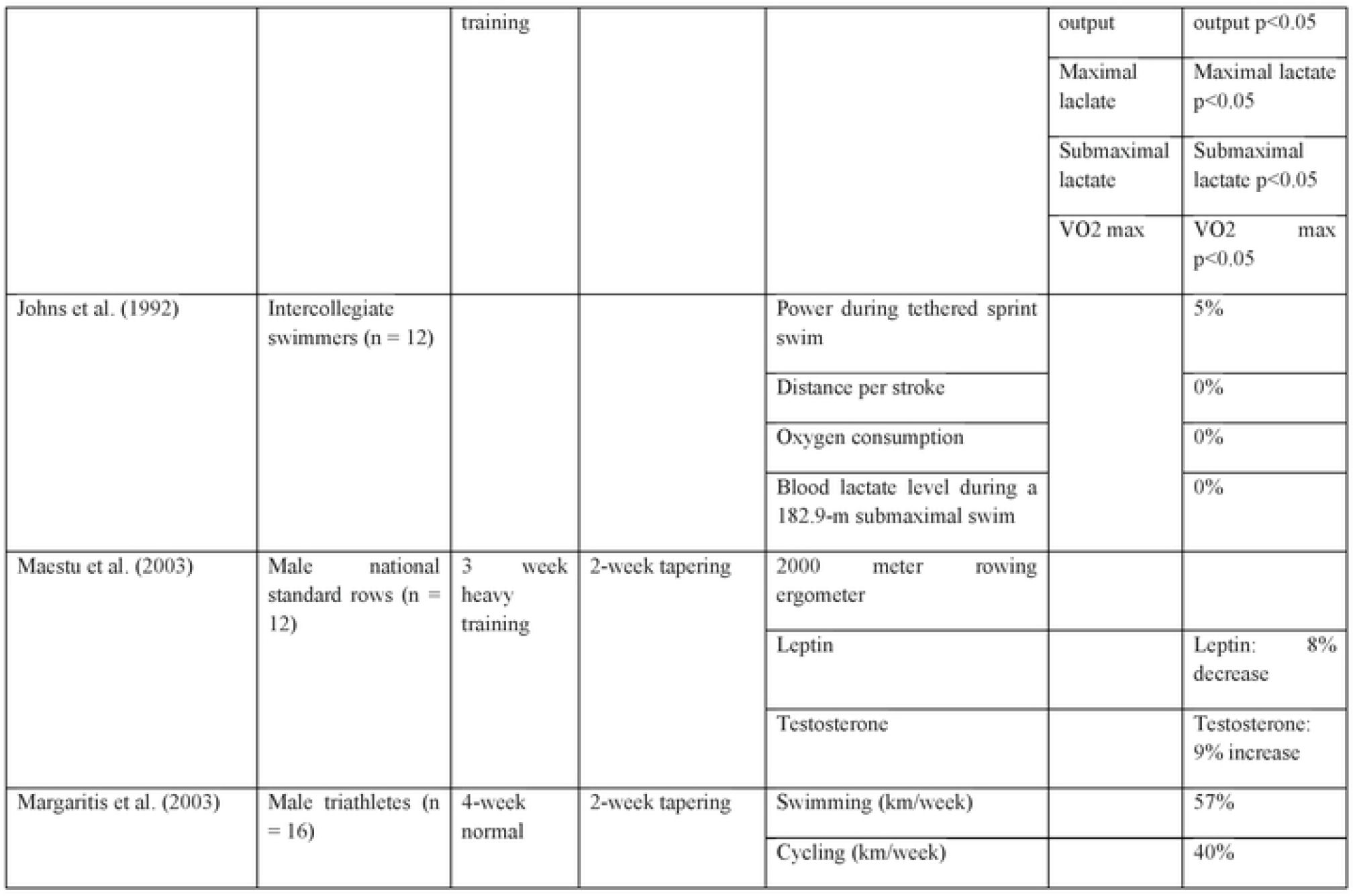
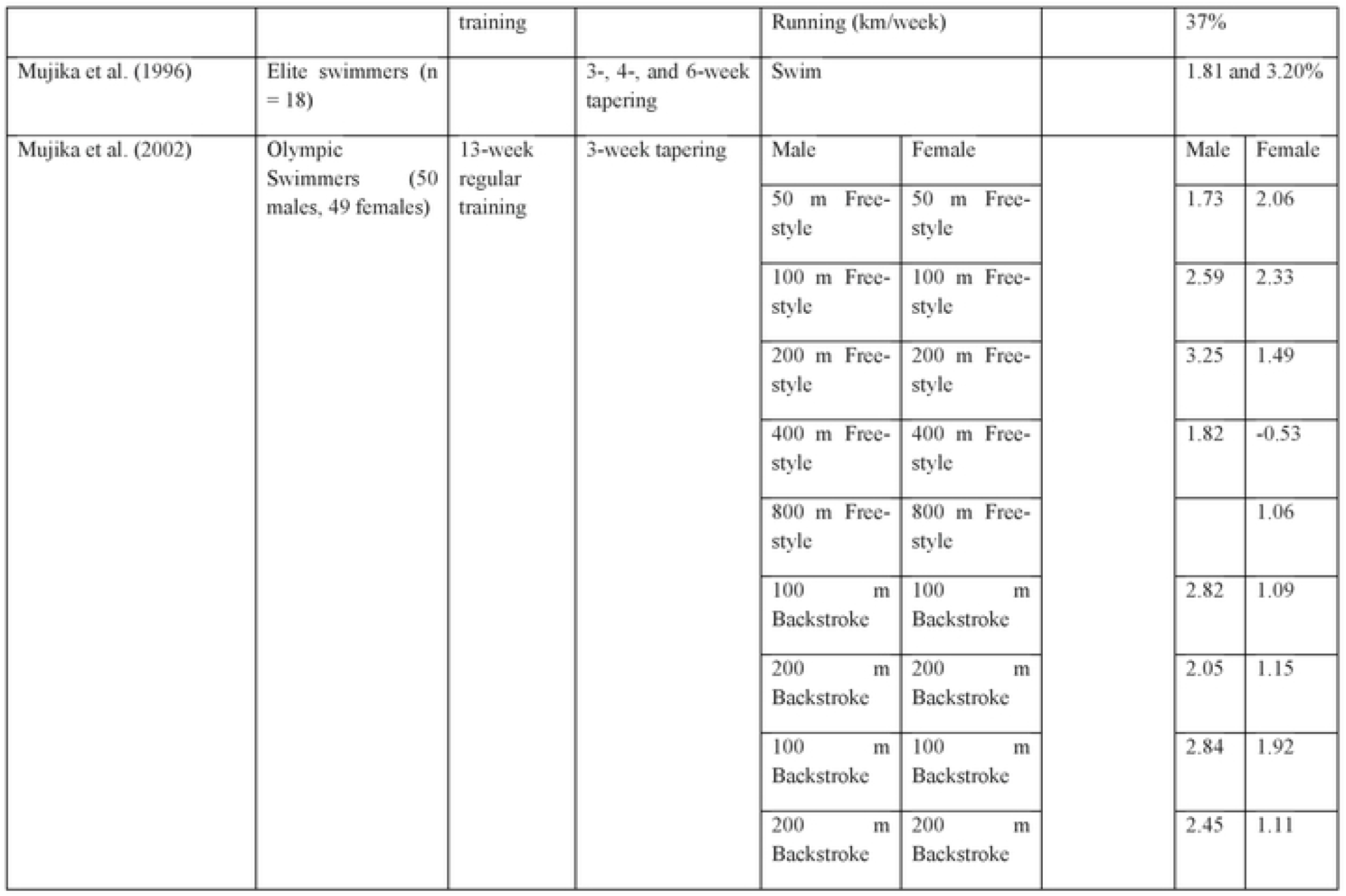
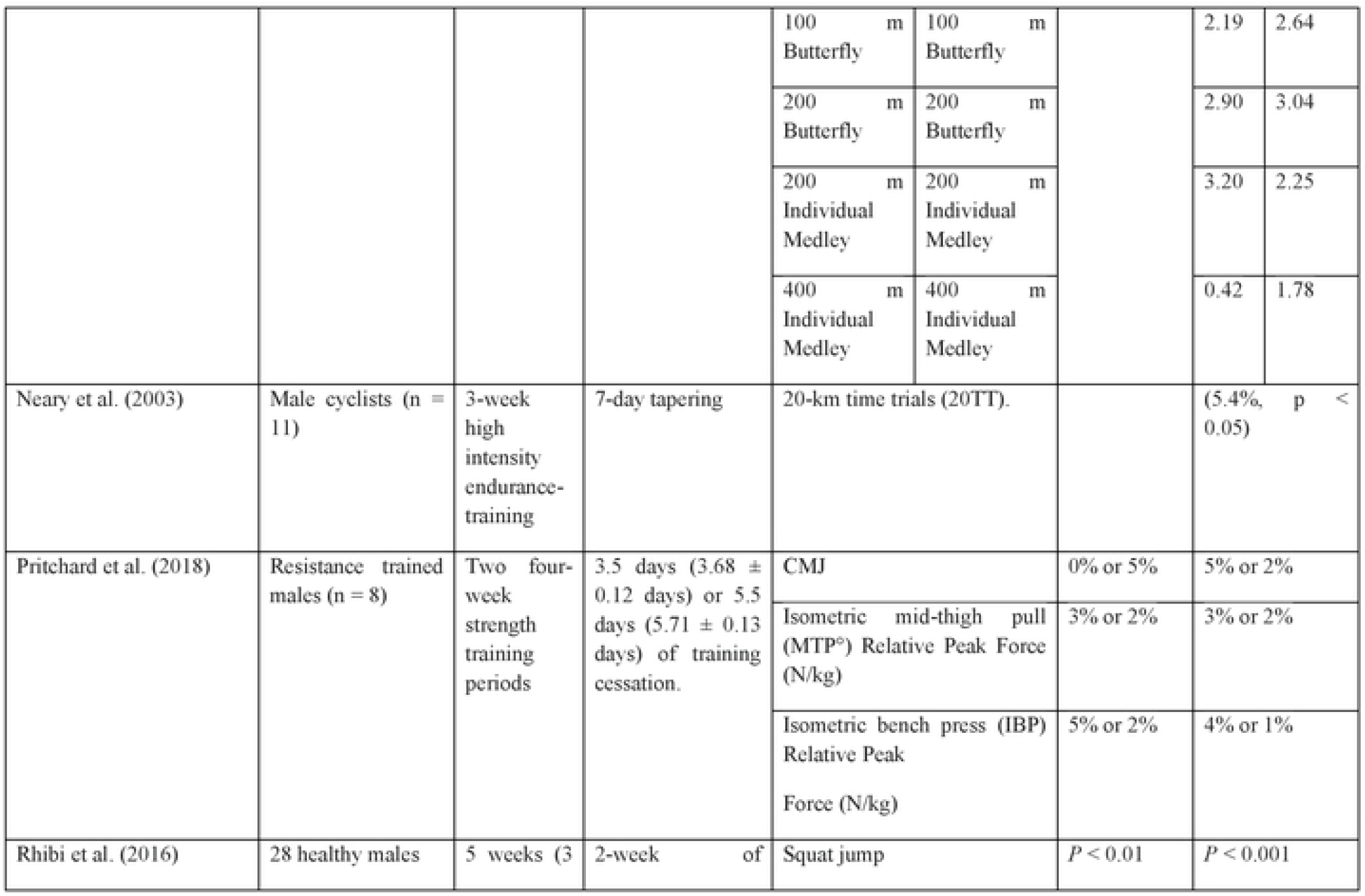
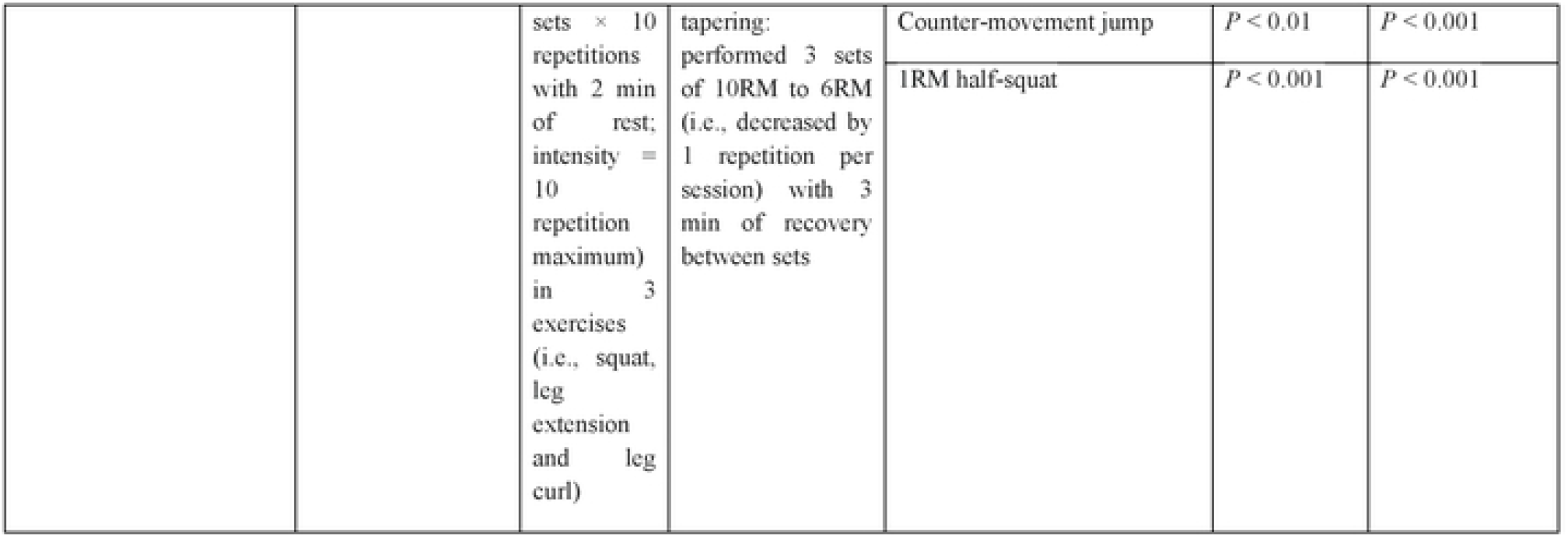
Effectiveness of various types of training and tapering phases on performance of different level and sport practice athletes.

| Author | Nature of sample | Training phase | Tapering phase | Test measure | Performance gain during training phase | Added performance gain during tapering |
| --- | --- | --- | --- | --- | --- | --- |
| de Lacey et al. (2014) | Professional rugby league players (age, $24.6 \pm 3.6$ years) | 4-month pre-season training period | Taper occurred during the final 21 days | Jump height | | 35% |
|  |  |  |  | Force-velocity profiles |  |  |
| | | | | $F_0$ | | 18% |
| | | | | $V_0$ | | 2.86% |
| | | | | $P_{max}$ | | 45% |
| Gibala et al. (1994) | Trained healthy Athletes males (Age, $23 \pm 2.1$ years) (n = 8) | 3-week | 10 days of reduced volume taper | Peak twitch | | 25% |
|  |  |  |  | Time to peak torque of elbow |  | 3% |
|  |  |  |  | Half-relaxation time of elbow |  | 0% |
|  |  |  |  | Maximum rate of torque development of elbow |  | 30% |
| Jeukendrup et al. (1992) | Male competitive cyclists (n = 8) | 2-week of intensified heavy | 2-week of recovery | Cycle ergometer test and outdoor 8.5 km time trial | Contests and maximal power | Contests and maximal power |
| | | training | | | output | output $p < 0.05$ |
| | | | | | Maximal lactate | Maximal lactate $p < 0.05$ |
| | | | | | Submaximal lactate | Submaximal lactate $p < 0.05$ |
| | | | | | VO2 max | VO2 max $p < 0.05$ |
| Johns et al. (1992) | Intercollegiate swimmers (n = 12) |  |  | Power during tethered sprint swim |  | 5% |
|  |  |  |  | Distance per stroke |  | 0% |
|  |  |  |  | Oxygen consumption |  | 0% |
|  |  |  |  | Blood lactate level during a 182.9-m submaximal swim |  | 0% |
| Maestu et al. (2003) | Male national standard rows (n = 12) | 3 week heavy training | 2-week tapering | 2000 meter rowing ergometer |  |  |
|  |  |  |  | Leptin |  | Leptin: 8% decrease |
|  |  |  |  | Testosterone |  | Testosterone: 9% increase |
| Margaritis et al. (2003) | Male triathletes (n = 16) | 4-week normal | 2-week tapering | Swimming (km/week) |  | 57% |
|  |  |  |  | Cycling (km/week) |  | 40% |

|  |  |  |  |  |  |  |  |  |
| --- | --- | --- | --- | --- | --- | --- | --- | --- |
|  |  | training |  | Running (km/week) |  |  | 37% |  |
| Mujika et al. (1996) | Elite swimmers (n = 18) |  | 3-, 4-, and 6-week tapering | Swim |  |  | 1.81 and 3.20% |  |
| Mujika et al. (2002) | Olympic Swimmers (50 males, 49 females) | 13-week regular training | 3-week tapering | Male | Female |  | Male | Female |
|  |  |  |  | 50 m Free-style | 50 m Free-style |  | 1.73 | 2.06 |
|  |  |  |  | 100 m Free-style | 100 m Free-style |  | 2.59 | 2.33 |
|  |  |  |  | 200 m Free-style | 200 m Free-style |  | 3.25 | 1.49 |
|  |  |  |  | 400 m Free-style | 400 m Free-style |  | 1.82 | -0.53 |
|  |  |  |  | 800 m Free-style | 800 m Free-style |  |  | 1.06 |
|  |  |  |  | 100 m Backstroke | 100 m Backstroke |  | 2.82 | 1.09 |
|  |  |  |  | 200 m Backstroke | 200 m Backstroke |  | 2.05 | 1.15 |
|  |  |  |  | 100 m Backstroke | 100 m Backstroke |  | 2.84 | 1.92 |
|  |  |  |  | 200 m Backstroke | 200 m Backstroke |  | 2.45 | 1.11 |
|  |  |  |  | 100 m<br>Butterfly | 100 m<br>Butterfly |  | 2.19 | 2.64 |
|  |  |  |  | 200 m<br>Butterfly | 200 m<br>Butterfly |  | 2.90 | 3.04 |
|  |  |  |  | 200 m<br>Individual<br>Medley | 200 m<br>Individual<br>Medley |  | 3.20 | 2.25 |
|  |  |  |  | 400 m<br>Individual<br>Medley | 400 m<br>Individual<br>Medley |  | 0.42 | 1.78 |
| Neary et al. (2003) | Male cyclists (n = 11) | 3-week high intensity endurance-training | 7-day tapering | 20-km time trials (20TT). |  |  | (5.4%, p < 0.05) |  |
| Pritchard et al. (2018) | Resistance trained males (n = 8) | Two four-week strength training periods | 3.5 days (3.68 ± 0.12 days) or 5.5 days (5.71 ± 0.13 days) of training cessation. | CMJ |  | 0% or 5% | 5% or 2% |  |
|  |  |  |  | Isometric mid-thigh pull (MTP°) Relative Peak Force (N/kg) |  | 3% or 2% | 3% or 2% |  |
|  |  |  |  | Isometric bench press (IBP) Relative Peak Force (N/kg) |  | 5% or 2% | 4% or 1% |  |
| Rhibi et al. (2016) | 28 healthy males | 5 weeks (3 | 2-week of | Squat jump |  | P < 0.01 | P < 0.001 |  |
| | | sets × 10 repetitions with 2 min of rest; intensity = 10 repetition maximum) in 3 exercises (i.e., squat, leg extension and leg curl) | tapering: performed 3 sets of 10RM to 6RM (i.e., decreased by 1 repetition per session) with 3 min of recovery between sets | Counter-movement jump | $P < 0.01$ | $P < 0.001$ |
| | | | | 1RM half-squat | $P < 0.001$ | $P < 0.001$ |

### Power and Maximal Strength

A few previous investigators have examined the effects of resistance training using a dynamic Olympic weightlifting exercises on the peak muscle power of handball players [18,22], but our study is the first to compare gains of peak power at weightlifting loads, using successive eccentric-concentric weightlifting exercises for the upper and lower body. The experimental subjects showed gains of absolute power for both the lower (18%; p<0.01) and upper (11%; p<0.01) extremities, although without significant change in relative power to body mass for the upper limbs. These results seem relatively in accordance with Arabatzi et al [29], who noted a significant increases in peak power output during the counter-movement jump, but not significant increases in peak power output during the squat jump, after an 8 week period (3 sessions per week) of Olympic weightlifting training in male physical education students. In contrast, Helland et al [30] saw no significant increases in peak power output during the CMJ, after 8 weeks period (3 sessions per week) Olympic weightlifting training, in football players.

There were further gains (p<0.05) in absolute power (W) of the lower limbs after tapering. De Lacey et al [27] also noted significant increases of power relative to body mass for the upper limbs (45%), as well as theoretical maximum force (18%), and the theoretical maximum velocity (2.86%) during the force-velocity test, after professional rugby player undertook a 21-day step taper.

Handball performance requires not only on strength, but also the ability to exert force at the necessary speed. We applied longer duration Olympic Weightlifting exercises with variable loads, judging that such a prescription was best for maximizing strength [29]. After the 10 weeks of resistance training, the experimental group out-performed the controls on all strength parameters, and this advantage persisted over the 2 weeks taper. Rhibi et al [31] also noted significant increases in 1-RM half-squat, after 12 weeks of lower-extremity resistance training followed by 2 weeks of tapering in health young men, and after 5 weeks of lower-extremity resistance training (p<0.01) followed by 2 weeks of tapering (p<0.01) in young volleyball players [31]. Likewise, Zaras et al [32] observed significant increases in 1-RM leg press of young and adult throwers, after 12 and 15 weeks of lower-extremity resistance training (p<0.05) followed by 2 weeks of tapering (p<0.05). Other studies of tapering have also seen significant increases in maximal strength, muscle power [33], and strength after 3-16 weeks of strength training [34]. Gibala et al [34] argued that 8 days of reduced training volume was sufficient to improve muscle strength. Likewise, Johns et al [13] reported 3% increases of muscle strength in swimmers after 10 and 14 days of tapering. Bosquet et al. [35] again suggested that two weeks of tapering was the optimal period to ameliorate physical performances and to eliminate accumulated fatigue. Loner periods of tapering seem undesirable because of the risk of detraining [36,37].

### Ball throwing velocity

After the initial training period, the experimental group showed greater velocities in all 2 types of ball throw (Table 2). Hermassi et al [22] also noted significant gains for all 3 types of ball throws following 8 weeks of heavy resistance training for both upper and lower limbs. Chelly et al [2018] noted gains with 8-weeks of plyometric training, and another report [38], described gains in elite male handball players from an 8-week resistance program. The present study seems the first to have demonstrated benefits from Olympic weightlifting exercises, and it underlines the benefits from 2 weeks of tapering. Others have shown the beneficial effect of 2 weeks of tapering on shot throws [32].

### Sprint performance and ability to change direction

Sprinting, rapid changes of direction, and acceleration are all important qualities in handball competition [22]. After the 10 weeks training period, a significant group x time interaction was found in 15m and 30m sprint performance (p≤0.001), and a further significant group x time interaction was observed for 15m performances after tapering. Tricoli et al [39] observed significant speeding of 10m sprint times in male physical education students, but no significant increases over 30 m, after 8 weeks of Olympic weightlifting training. Others [40] observed significant increases in 25m sprint times in male collegiate athletes after 12 weeks of Olympic weightlifting training, and enhancement of Ayers et al [41] observed significant increases in 40-yard but not 30 m sprint times of female collegiate athletes after 6 weeks of Olympic weightlifting training, although Helland et al [30] saw no significant improvements in 40-yard sprint times after 8 weeks of Olympic weightlifting in football players, and Hoffman et al [44] saw no improvements in 30 m sprint time after young athletes underwent 15 weeks of Olympic weightlifting.

This is the first investigation to have studied the effects of tapering on ball throwing velocity, but others have studied the effect of tapering on repeated-sprint performance; Bishop et al [43] observed no-significant increase of total work (4.4%, p = 0.16) or peak power (3.2%; p = 0.18) in female athletes during the 5 × 6-s test, but they did see a lessening of work decrement (7.9 ±4.3% decrease; p<0.05) and a significant increases in shot throw performance, after 10 days of tapering. Other authors have shown both increases and decreases in change of direction performance [44-48]. The current investigation seems the first to have studied the effects of tapering on change of direction ability; we saw a small reduction of ability on the T-half test (−0.7%) after tapering.

## PRACTICAL APPLICATIONS

Short-term resistance training using weightlifting exercises offers a stimulus that is uniquely different from power lifts, and should be a component of any resistance training program for handball players, who require quick, powerful movements. Tapering reduced cumulative fatigue and further increased maximal strength, vertical jump, and ball velocity performances. Coaches may prefer to use hierarchical resistance lifts programs when there is a need to improve power, strength, sprint, ability to change direction and throwing abilities, since all of these abilities are enhanced by this type of training. Alternatively, resistance training could precede Olympic weightlifting training, so that participants can first achieve an increase of muscle strength and joint stability, allowing them to perform power specific exercises to enhance their performance. Most of the gains associated with tapering seem of substantial size, and should thus be of interest for both handball players and their coaches. We would also encourage further investigation of the many potential factors underlying the increased performance during tapering.

## Acknowledgments

This work was supported by the Sport Science Program, College of Arts and Sciences, Qatar University, Doha, Qatar.

